# UNI-EM: An Environment for Deep Neural Network-Based Automated Segmentation of Neuronal Electron Microscopic Images

**DOI:** 10.1101/607366

**Authors:** Hidetoshi Urakubo, Torsten Bullmann, Yoshiyuki Kubota, Shigeyuki Oba, Shin Ishii

**Author notes:** Correspondence: Hidetoshi Urakubo.

## Abstract

Recently, there has been a rapid expansion in the field of micro-connectomics, which targets the three-dimensional (3D) reconstruction of neuronal networks from a stack of two-dimensional (2D) electron microscopic (EM) images. The spatial scale of the 3D reconstruction grows rapidly owing to deep neural networks (DNNs) that enable automated image segmentation. Several research teams have developed their own software pipelines for DNN-based segmentation. However, the complexity of such pipelines makes their use difficult even for computer experts and impossible for non-experts. In this study, we developed a new software program, called UNI-EM, that enables 2D- and 3D-DNN-based segmentation for non-computer experts. UNI-EM is a software collection for DNN-based EM image segmentation, including ground truth generation, training, inference, postprocessing, proofreading, and visualization. UNI-EM comes with a set of 2D DNNs, i.e., U-Net, ResNet, HighwayNet, and DenseNet. We further wrapped flood-filling networks (FFNs) as a representative 3D DNN-based neuron segmentation algorithm. The 2D- and 3D-DNNs are known to show state-of-the-art level segmentation performance. We then provided two-example workflows: mitochondria segmentation using a 2D DNN as well as neuron segmentation using FFNs. Following these example workflows, users can benefit from DNN-based segmentation without any knowledge of Python programming or DNN frameworks.

## Introduction

In recent years, there has been a rapid expansion in the field of micro-connectomics, which targets the three-dimensional (3D) reconstruction of neuronal networks from a stack of two-dimensional (2D) electron microscopic (EM) images^1-3^. Neuroscientists have successfully reconstructed large-scale neural circuits from species, such as mouse^4^, Drosophila melanogaster^5^, and Zebrafish^6^. Neuronal boundary detection (neuron segmentation) of large numbers of EM images are required for such large-scale reconstructions, and its automation is critical even for smaller-scale segmentation.

To facilitate automated segmentation, neuroscientists organized the EM Segmentation Challenge in the International Symposium on Biomedical Imaging 2012 Conference (ISBI 2012)^7,8^. The benchmark studies have demonstrated the effectiveness of a specific class of deep neural networks (DNNs), i.e., convolutional neural networks (CNNs) (e.g., Ciresan et al.^9^). Especially, U-Net had a large impact owing to its highest EM segmentation accuracy and wide applicability to semantic labeling^10^. Further, public 3D EM datasets such as FIB-25 (Drosophila)^11^, SEGEM (mouse cortex)^12^, and 3D segmentation of neurites in EM images (SNEMI3D, mouse cortex) have been extensively used for the benchmarks of 3D CNNs^13^. Januszewski et al. developed a recursive 3D CNN, called flood filling networks (FFNs), which showed the best segmentation accuracy in FIB-25 and the second-best accuracy in SNEMI3D^14^. In many cases, source codes of these CNNs are available at public repositories. However, EM segmentation relies not only on CNNs, but also on other procedures, such as ground truth generation, pre/postprocessing, proofreading, and visualization. Thus, these CNNs should work as a part of system software.

Advanced connectomics laboratories have developed their own software pipelines to obtain benefits from CNN-based segmentation including Rhoana^15,16^, Eyewire^17^, NeuTu^18^, and DVID^19^. The main objective for these pipelines is large scale 3D reconstructions conducted by large teams including computer experts for setup and maintenance. They are too complex for smaller teams. EM segmentation has also been targeted by many sophisticated standalone software packages, such as Reconstruct^20^, Ilastik^21^, Knossos^22^, Microscopy Image Browser^23^, and VAST^24^. Unfortunately, none of these software packages incorporates DNN-based segmentation. A recent attempt to include the power of DNNs in biomedical image processing is an U-Net plug-in for the widely used ImageJ^25^. ImageJ can run on any Java virtual machine (JVM) regardless of the underlying operating system. However, for DNNs, the most supported programming language is not Java, but Python. Although ImageJ is extensible via plugins written in Java, the implementation of new DNN models is still technically demanding.

We thus developed a unified Python-based software to conduct DNN-based automated EM segmentation (UNI-EM) aimed at non-computer experts. UNI-EM utilizes the widely used Tensorflow framework^26^ for the implementation of several 2D CNNs^10,27-29^ and 3D FFNs^14^. Furthermore, it includes a series of 2D/3D filters for classic image processing as well as the proofreading software Dojo^30^. Those features enable users to follow the procedure of DNN-based segmentation, i.e., ground truth generation, training, inference, postprocessing, proofreading, and visualization. The users do not need to install Python and any modules, because we also provide Python installation-free versions of UNI-EM (pyinstaller version). Together, using UNI-EM, non-computer experts can benefit from DNN-based segmentation on the most widely used operating system (OS) for desktop computers, Microsoft Windows 10 ^31^.

## Results

### Outline of software

UNI-EM is a software collection for DNN-based EM image segmentation that includes ground truth generation, training, inference, postprocessing, proofreading, and visualization (Fig. 1). UNI-EM is written in Python 3.6 and runs on Microsoft Windows 10 (64 bit). We also built UNI-EM on the Python application bundler “pyinstaller,” thus users can use UNI-EM without installing the Python programming environment. CPU and GPU versions are available, and users can maximize the performance using the GPU version if the computer is equipped with an NVIDIA GPU card that has >3.5 NVIDIA compute capability. Next, we introduce the components of UNI-EM.

**Figure 1.**
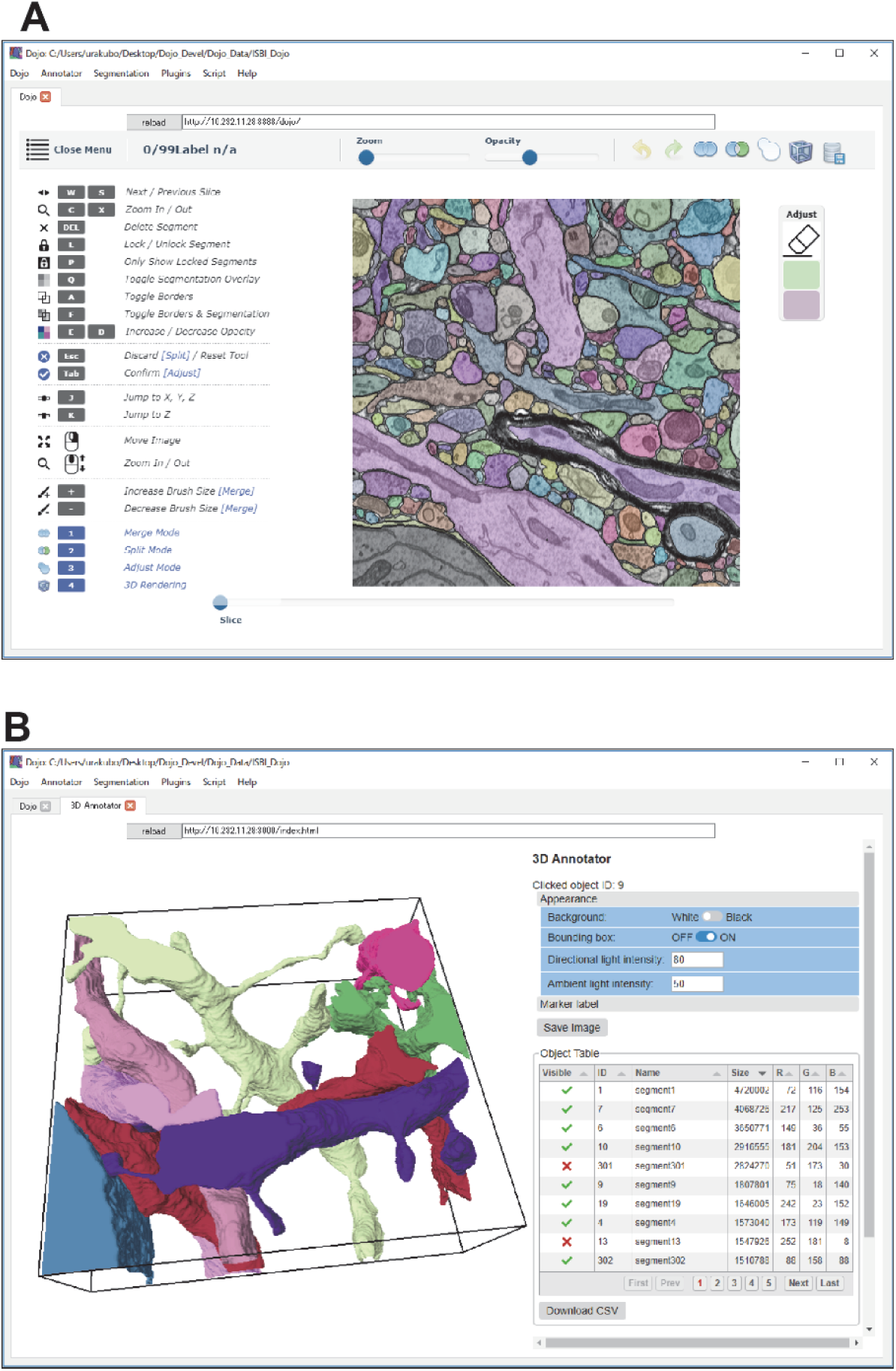
GUIs of UNI-EM. **A.** Proofreader Dojo with extension. The GUI of Dojo was reorganized. Users can correct mis-segmentation as well as make ground truth using paint functions. The reorganized Dojo supports import/export functions of EM/segmentation image stack files. **B.** A 3D annotator. A 3D viewer (left) is associated with the object tables (right) that display segmented object and marker points. Visualization results and tables are exportable as png and csv files, respectively. The GUIs in **A** and **B** are provided as web applications. Multiple users can access these GUIs through equipped or external web browsers.

The main component of UNI-EM is a web-based proofreading software, Dojo (Fig. 1A)^30^. Dojo provides a graphical user interface (GUI) for users to correct mis-segmentation arising from automated EM segmentation. We extended Dojo to have file import/export functions (png/tiff files), a more sophisticated GUI, and multiscale paint functions. With these extensions, users can use Dojo not only for proofreading, but also for ground truth generation, both of which are important manual operation procedures for DNN-based segmentation. Dojo consists of a Python-based web server and an HTML5/JavaScript-based client interface. The server-client system allows multiple users to simultaneously access it through web browsers in an OS-independent manner. UNI-EM equips its own web browser “chromium” for the standalone use of Dojo with either a mouse or a stylus.

We also developed a new 3D annotator to visualize the proofread objects in a 3D space as well as to annotate these segmented objects (Fig. 1B). This is a surface mesh-based 3D viewer with a table that shows segmented objects. Users can change the color and the brightness of target objects, and export the visualization results as a png image file. Users can also assign a name to each object, and put marker points on the object surface. The results of these annotations can be exported as csv files for further analyses.

We then implemented a U-Net equipped with a GUI as a representative 2D CNN for EM-image segmentation^10^. U-Net has a characteristic contracting and expansive convolution layers with skip connections, which generated the highest segmentation accuracy in ISBI 2012 at the time of publication^10^. The original form of U-Net was implemented with expandable convolution/deconvolution (transposed convolution) layers. We similarly implemented ResNet, which was the winner of the ImageNet Large Scale Visual Recognition Challenge (ILSVRC) 2015, as well as the winner of Microsoft COCO 2015 detection and segmentation^27^. Highway-Net^28^ and Dense-Net^29^ were also implemented. Users can choose any combination of these CNNs, loss functions, and training times through a command panel. The command panel enables 2D CNN-based segmentation without any knowledge of Python programming language.

We further wrapped FFNs as a representative algorithm of 3D CNN-based neuron segmentation^14^. FFNs are a recurrent CNN that infers a volume mask indicating whether target voxels belong to the centered object, and the inference program obtains an overall volume mask for each object using a flood filling algorithm. FFN outperformed many other algorithms in the segmentation accuracies of FIB-25 and SNEMI3D, and FFN is the best in publicly available algorithms. We equipped FFNs with a command panel so that users can conduct the preprocessing, training, inference, and postprocessing through the command panel.

The 2D CNNs and 3D FFNs were implemented on Tensorflow framework^26^. Its resource monitor Tensorboard can be conveniently accessed from UNI-EM. By selecting “Segment → Tensorboard” from the dropdown menu, users can easily check the status of a target CNN, such as network topology and loss function. Also, users can implement their own DNN algorithms at the “Plugin” dropdown menu. Details on how to implement a new DNN are outlined in an online manual. In addition, UNI-EM has a GUI for 2D/3D classic image filters. Users can apply multiple image filters simultaneously to a stack of 2D images by a single execution. All the source code with the online manual is provided at https://github.com/urakubo/UNI-EM.

### Example workflows

In this section, we demonstrate how users can benefit from UNI-EM by introducing two example workflows. The first one is the mitochondria segmentation using 2D CNNs, and the second one is the neuron segmentation using 3D FFNs, which is required by micro-connectomics.

In both cases, we targeted an EM-image stack that was prepared for SNEMI3D^32^. The target brain region is the mouse somatosensory cortex, and the EM images were obtained using a scanning electron microscopy (SEM) in combination with an automatic tape-collecting ultra-microtome system (ATUM/SEM)^33^. The spatial resolution of the EM images is 6 nm per pixel (xy-plane) and 30 nm per Z slice, and the overall image volume is 6.1 × 6.1 × 3 μm. They were passed through the contrast-limited adaptive histogram equalization filter (CLAHE; block size 127, histogram bins 256, max slope 1.50) before segmentation.

#### Case 1: Mitochondria segmentation using 2D CNN

Mitochondria are abundant where the metabolic demand is high, such as synapses and active axons^34,35^, and they possess very characteristic shapes^36^. Their detection and quantification are important for treating neuronal diseases^37^. Importantly, 2D CNNs yield sufficient segmentation accuracies^38,39^.

Although mitochondria segmentation is a good target for 2D CNN-based segmentation, it was not accessible to inexperienced users (Fig. 2A). First, the inexperience users need to learn how to use Python, install a DNN frame work, and download an implementation of a target CNN from a public repository. The other software packages need to be installed for ground truth generation, post-processing, and proofreading (Fig. 2A). These steps might be learned, but a major hurdle is the transfer of data especially to a CNN, when the users must convert EM/segmentation images to HDF5 or npz format files. We confirmed that UNI-EM successfully decreases those demanding tasks as follows (Fig. 2B). Two test users (H.K. and Y.F.) without knowledge of Python programming conducted the following procedure (Fig. 2C):

**Figure 2.**
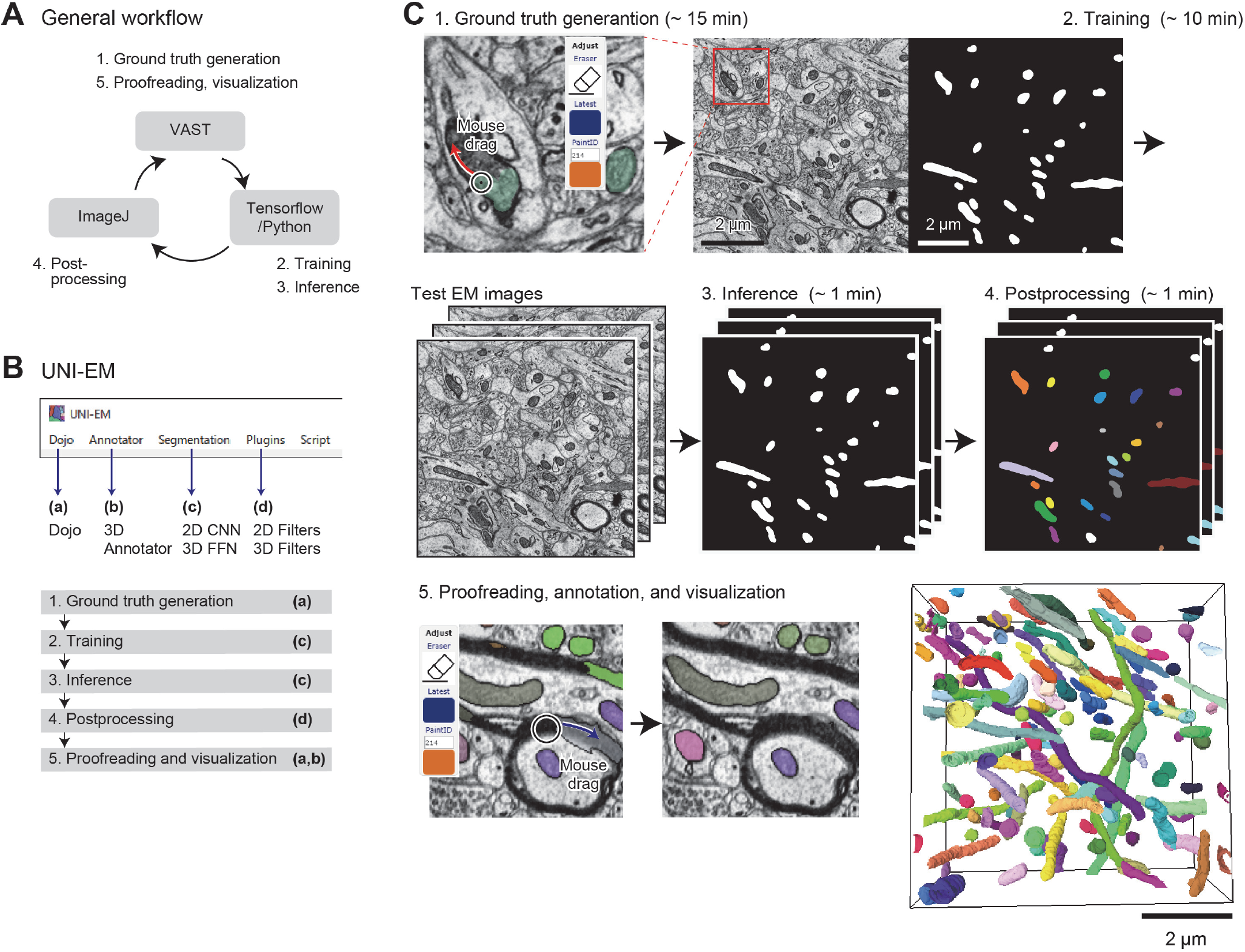
Example workflow 1: Mitochondria segmentation using 2D CNN. **A.** General workflow. Users first paint mitochondria regions of a target EM image with painting software, e.g., VAST (1, top) ^24^. This mitochondrial segmentation image (ground truth) and the EM image are transferred to Tensorflow/Python for DNN training and inference (2,3; right). Inferred segmentation is postprocessed (4, left), e.g., using imageJ, then proofread and visualized by VAST (5, top). Such relays between software packages are necessary. **B.** UNI-EM dropdown menu. It locates a series of software (a-d) for the procedure of DNN-based segmentation (1-5). Standard png/tiff file format is used to connect these software packages. **C.** Workflow in UNI-EM. Extended Dojo supports paint functions (1; top, left) to draw mitochondrial segmentation (top, right). Users can conduct DNN training (2) and inference (3) through a control panel. Labeling function is also implemented for postprocessing (4, each label is denoted by color). These segmented images are proofread by Dojo (5, left), and visualized by a 3D annotator (5, right).

1. Ground truth generation. The test users were requested to paint mitochondrial regions of a single EM image using UNI-EM (Dojo). The generated ground truth was exported as an 8-bit grayscale PNG file (− 20 min).
2. Training. A 16-layer ResNet was trained with the ground truth (−10 min computation time).
3. Inference. The trained ResNet was applied to test the EM images to obtain inferred 2D segmentation (−1 min).
4. Postprocessing. The inferred 2D segmentation images were binarized, and then each isolated region in 3D space was labeled with a specific ID number (−10 min).
5. Proofreading, annotation, and visualization. The test users proofread it with Dojo and visualized it with the 3D annotator (−30 min).

The test users successfully conducted the above procedure within the time indicated in parentheses, and the segmentation accuracy was high (Fig. 2C, 5; RAND score: 0.85; see Methods), as expected from published results of CNN-based segmentation^38,39^. The detailed instruction for the mitochondria segmentation task can be found at https://github.com/urakubo/UNI-EM.

In this process, we requested the test users to use a 16-layer ResNet for mitochondrial segmentation. This request was determined based on the following quantitative survey on the segmentation accuracies of mitochondria, synapses, and neurons (Fig. 3A). Here we utilized the RAND score as a measure of segmentation accuracy. The larger RAND score denotes higher accuracy. We first confirmed that only one ground truth image was sufficient for the segmentation of mitochondria (Fig. 3B), and 10 ground truth images are sufficient for neurons and synaptic segmentations. We then confirmed that the square, dice, and logistic loss functions were appropriate for segmentations (Fig. 3C). All the types of 2D CNNs showed high accuracies in mitochondria segmentation (Fig 3D, green lines; > 0.9 RAND score). In addition, U-Net was not appropriate for membrane segmentation (Fig. 3D, red line; ∼ 0.3 RAND score), and the segmentation accuracies in synapses are not high regardless of the types of CNNs (∼ 0.3 RAND score, Fig. 3D). The accuracy of mitochondria segmentation in the standard CNN (network topology: ResNet;, loss function: least square, number of layers, : 9, training epochs: 2000; number of training images: 5) was indeed comparable with the accuracy in a recent 3D CNN-based, state-of-the-art algorithm^39^. Their segmentation accuracy was quantified as Jaccard 0.92, Dice 0.96, and conformity 0.91 (ATUM/SEM data), whereas the segmentation accuracy of our standard CNN was Jaccard 0.96, Dice 0.93, conformity 0.93. Here, the metrics, Jaccard, Dice, and conformity, show segmentation accuracy^39^, and their larger scores indicate higher accuracy. Further, their 3D CNN requires 77 h train time with NVIDIA K40 GPU, whereas our standard CNN required only 10 min under a similar condition. In addition, the 3D CNN is trained with 3D ground truth, which requires a large amount of tedious manual labeling. Overall, the implemented 2D CNN-based segmentations showed sufficiently high and competitive accuracies compared to the current state-of-the-art mitochondrial segmentation algorithms^39^.

**Figure 3.**
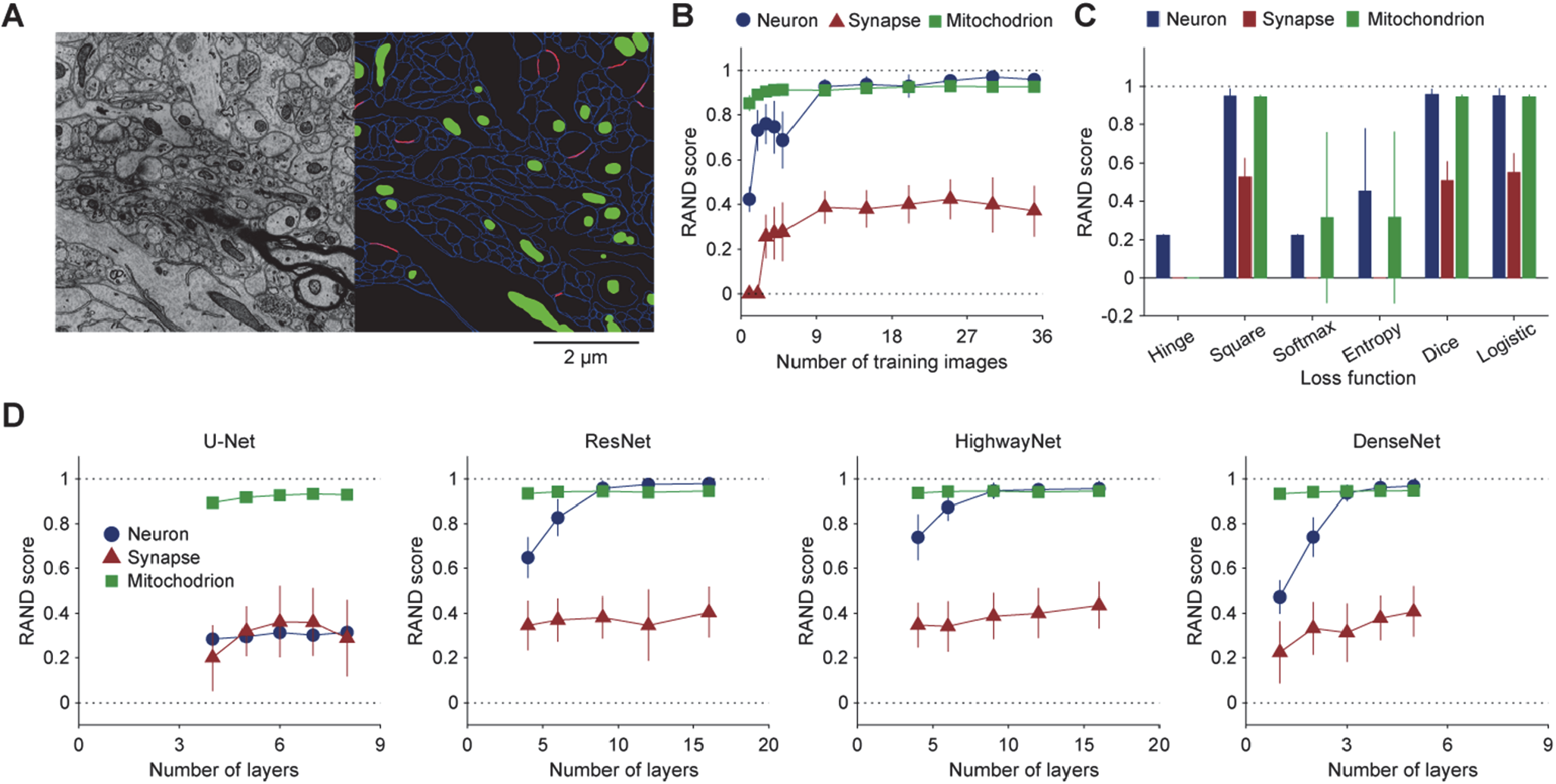
Performance survey in 2D CNN-based segmentation of neurons, synapses, and mitochondria. **A.** A part of target EM images (left, SNEMI3D) ^11^ and ground truth segmentation (right). Each image panel has 1024 × 1024 voxels (3 nm/voxel in x-y plane), and 100 z-slices (3 nm/voxel in z slice). In the right panel, blue and lines indicate cellular membranes and synapses, respectively, and green areas indicate mitochondria. **B.** Training image number dependence of segmentation accuracy (n = 15, mean ± SD; RAND score, see Methods). The RAND score becomes close to 1 if the inferred segmentation is similar to ground truth. **C.** Loss function dependence of segmentation accuracy (n = 60, mean ± SD). Here, ‘Square’ denotes least square, ‘Softmax’ denotes softmax cross entropy, and ‘Entropy’ denotes multi-class multi-label cross entropy. **D.** Network topology dependence of segmentation accuracy (n = 15, mean ± SD). In B-D, all the parameters except the target parameters were set as follows: the number of training images: 1; loss function: least square; network topology: ResNet; number of layers: 9; training epochs: 2000; number of training images: 5 (the standard CNN). The 2000 training epochs gave steady states of their losses.

#### Case 2: Neuron segmentation using 3D FFN

We next asked a test user (N.Y.) to conduct neuron segmentation using 3D FFNs^14^, which is a main topic of micro-connectomics. A variety of 2D and 3D CNNs have been proposed for accurate neuron segmentation^9,13^. FFNs currently show one of the highest segmentation accuracies in neuron segmentation, although they require laborious work to generate 3D ground truth. Users can certainly generate 3D ground truth with Dojo, although we recommend VAST lite for drawing 3D ground truth^24^. In our test case, we used the ground truth included in the SNEMI3D dataset. The test user successfully conducted the following procedure through the command panel (Fig. 4A):

**Figure 4.**
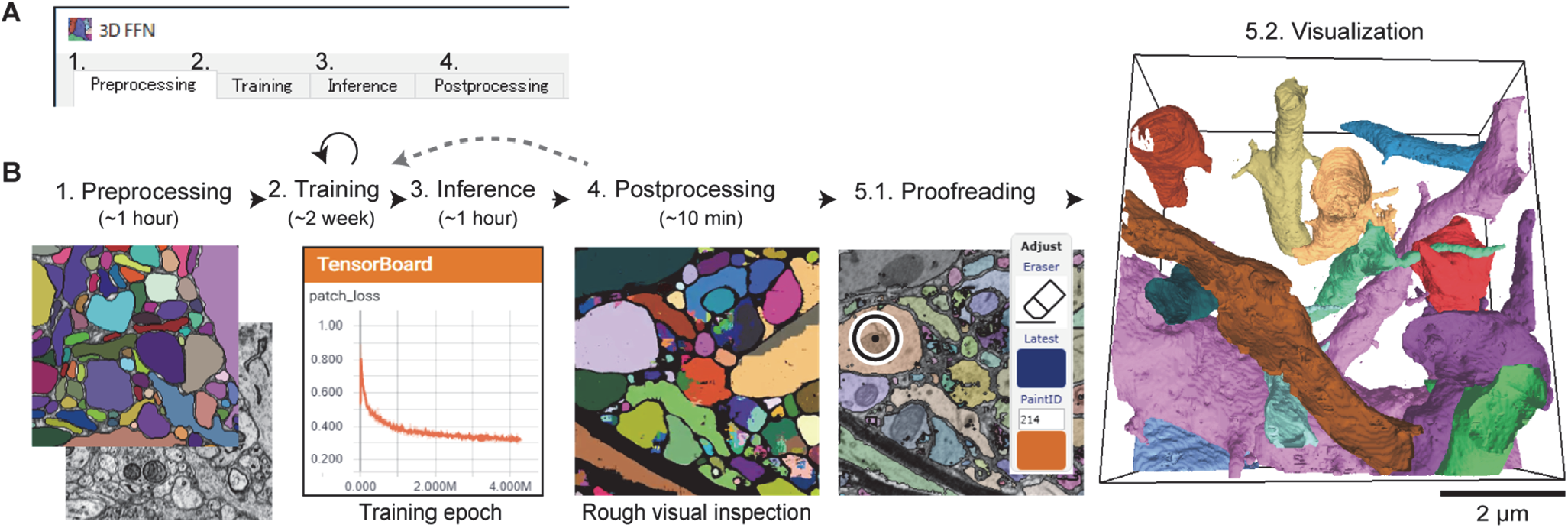
Example workflow 2: Neuron segmentation using 3D FFN. **A.** Control panel of 3D FFNs. Each tab (1-4) has one execute button for each FFN process. **B.** Workflow. Computation times are indicated in parentheses. (1) Preprocessing. Ground truth segmentation and EM images are converted to intermediate files. (2) Training. FFNs are trained with the intermediate files. Users can monitor the progress of training using Tensorboard. (3) Inference. (4) Postprocessing. The program can also generate colored inferred segmentation for rough visual inspection. If the segmentation quality is insufficient, users can further continue the training process. (5.1) Proofreading using Dojo. (5.2) Visualization by a 3D annotator.

1. Preprocessing. Stacks of target EM images and ground truth images were converted to FFN-specialized style files (−1 h computation time; Fig. 4B).
2. Training. The FFNs were trained with the preprocessed EM-image/segmentation files (−2 weeks computation time with a NIVDIA GTX1080ti GPU; Fig. 4B).
3. Inference. The trained FFNs were applied to a stack of test EM images for the inference of 3D segmentation (−1 h computation time with a NIVDIA GTX1080ti GPU; Fig. 4B).
4. Postprocessing. The output segmentation files were converted to a PNG file stack (−10 min computation time; Fig. 4B).
5. Proofreading and visualization. The converted PNG files and EM images were imported to Dojo for proofreading as well as the 3D annotator for visualization (Fig. 4B).

Note that trained FFNs directly infer a 3D segmentation from a stack of 2D EM images. FFNs gave a reasonable neuron segmentation (Fig. 4B right), and its segmentation accuracy was 0.84 (RAND score, 7 million training epoch; see Methods)^14^. This score was obtained without any postprocessing and specific parameter turning for the SNEMI3D dataset, and the topological structure of neurites were fairly preserved in the segmentation results. Januszewski et al. have reported the 0.975 RAND score in the case of the SNEMI3D dataset^14^. This score was obtained with the two additional processes: the automated agglomeration of oversegmentation and a 2D watershed^14^. Thus, there is a room for further improvements. Although FFNs require very long training time (− two weeks), users can benefit from its precise inference that drastically decreases tedious proofreading work.

### System design

UNI-EM is developed under the Python development environment and the Python bindings for v5 of the Qt application framework for GUI (PyQt5). The combination of Python and PyQt5 is typical for Python GUI desktop applications (e.g., Sommer et al.^21^), and UNI-EM utilizes this combination for GUI-equipped 2D CNNs and 3D FFNs (Fig. 5). The desktop application style is appropriate for DNN because DNN training/inference generally occupies all GPU resources of a desktop computer, and the shared use of GPU is ineffective. On the other hand, Dojo, a 3D annotator, and Tensorboard are web applications. The web application style provides remote accessibility to these applications, hence multiple users can simultaneously use them (remote users in Fig. 5). Tensorboard enables the remote observation of progression in DNN training. Dojo enables multiple users to correct mis-segmentation simultaneously, and the 3D annotator enables multiuser annotation. Together, UNI-EM is comprised of desktop and web application systems, and this heterogeneity improves the usability and effectiveness of this software in real-world applications.

**Figure 5.**
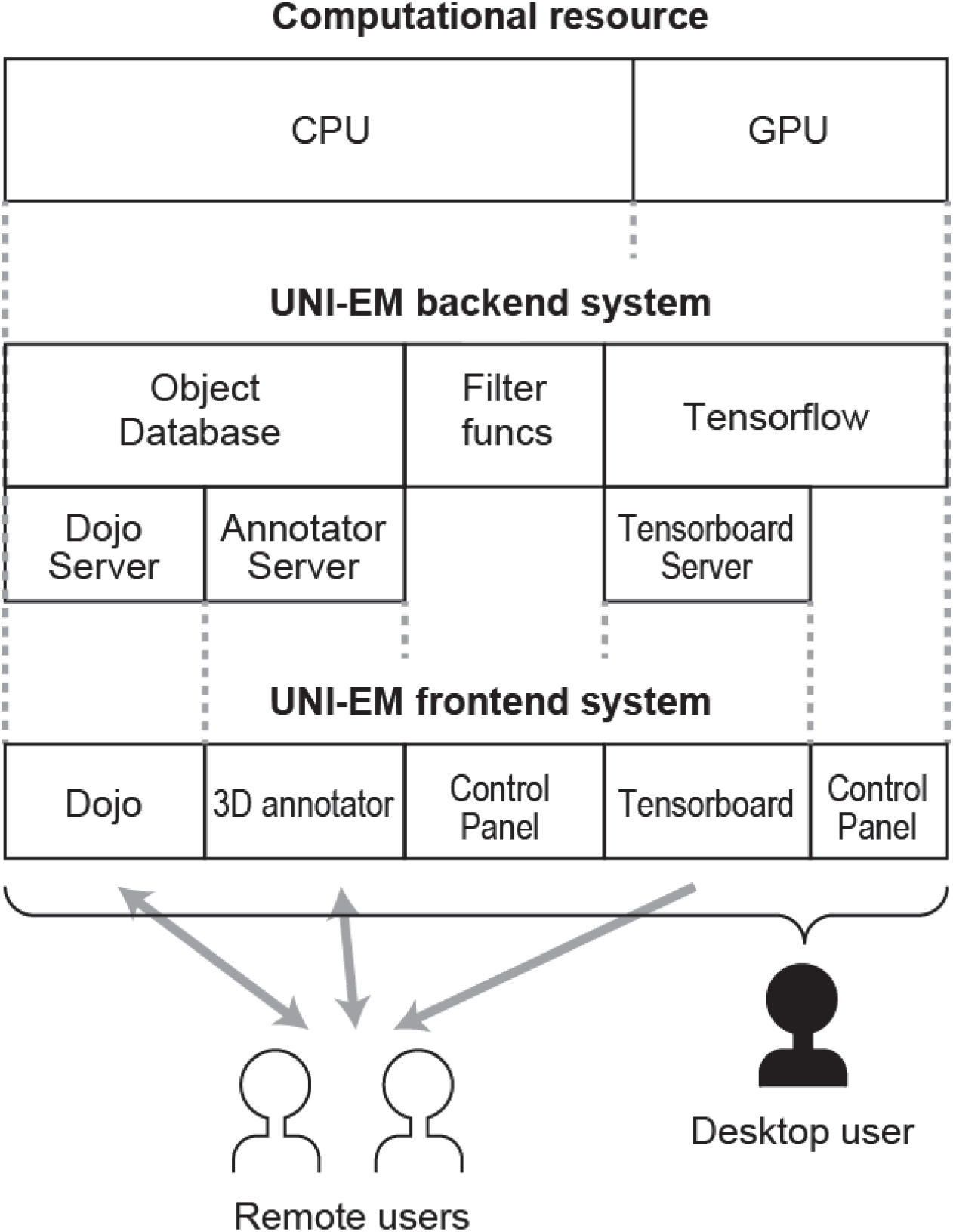
Underlying architecture of UNI-EM. UNI-EM is a heterogenous system. Present desktop computers have two types of computational resources: CPU and GPU (top). GPU is used by Tensorflow for DNN computing (middle), which is not appropriate for shared use. Only its resource monitor Tensorboard can be used by remote users (bottom). Similarly, remote users can use proofreader Dojo and 3D annotator. Only a desktop user (silhouette person) can control all UNI-EM functions including a job submission for DNN computing such as training and inference.

## DISCUSSION

We present UNI-EM as a software package for DNN-based automated EM segmentation. UNI-EM unifies pieces of the software for DNN-based segmentation. We validated its effectiveness in two example workflows: Mitochondria segmentation using a 2D CNN and neuron segmentation using 3D FFNs. Users without any knowledge of Python programming can follow the overall processes, and their segmentation accuracies are comparable with state-of-the art methods. Thus, UNI-EM is a useful tool for non-computer experts.

In recent years, DNNs has rapidly become popular in generic image segmentation as well as EM image segmentation^8^. Many DNN-based segmentation algorithms have been proposed, and in many cases their source codes are released together with the scientific publication. Nevertheless, it is still difficult for non-computer experts to test such DNN algorithms for their own EM images. They are located at public repositories where they are sometimes hard to find, and many algorithms require their own format input/output files, which require the knowledge of Python or other programming languages. In the current version, UNI-EM bundles only a few DNN algorithms, but it can easily incorporate such new DNN algorithms.

Two-dimensional CNN-based segmentation with subsequent Z-slice connection into three-dimensional objects is one of the best choices if the target objects do not have complicated shapes like neurons. In the example workflow, the test users successfully extracted the oval-shaped mitochondria within 2 h, and the segmentation accuracy was higher than those of conventional machine learning methods such as AdaBoost^15^. Two-dimensional CNNs can also be applied to neuron segmentation, but their accuracies have been surpassed by 3D CNNs^13,14,40,41^. As one of such 3D CNNs, FFNs yield very accurate neuron segmentation. Nevertheless, there are at least two remaining barriers for its common use. First, FFNs require very long training times (over one week). Second, they require a certain amount of 3D ground truth segmentation. In our experience, two-week labor was required to manually draw 3D ground truth even using a sophisticated paint tool^24^. FFNs are of course still an excellent selection if we consider the time for manual correction of mis-segmentation arising from other segmentation methods.

The main component of UNI-EM is the proofreading software Dojo with extensions^30^. Similar to Dojo, many excellent proofreading/manual annotation software programs are available, e.g., Reconstruct^20^, Ilastik^21^, TrakEM2 ^42^, VAST^24^, Knossos^22^, Microscopy Image Browser^23^, CATMAID ^43^, and NeuTu^18^. The primary advantage of Dojo is its architecture. It is designed as web application and can serve for both standalone or collaborative uses. Some proofreading software utilizes the vectorized outline representation of segmentation that can work with less memory consumption^20^. The voxel-based representation utilized by Dojo is more memory-intensive. However, almost all DNN-based algorithms produce voxel-based segmentation; thus, voxel-based proofreaders such as Dojo are necessary for correcting their mis-segmentation.

Web application systems are widely utilized in many types of software^43,44^. This design principle has many advantages; i.e., there is no need for the end users to install any software except for a web browser, OS independency, cloud resource accessibility, and it usually includes multiuser access. It also has drawbacks compared with a desktop application system when server computation resources or network bandwidth are limited. UNI-EM takes advantages from both of them by utilizing a heterogeneous desktop/web application system. Dojo web application provides multiuser accessibility, and its backend server runs on a Python environment. The Python desktop system allows the intensive use of GPU computing and the tile-based pyramidal data structure stored in a hard disk drive. The desktop system does not have the limitation of the transfer of large amounts of data to cloud systems.

Note that almost all the programs of UNI-EM are written in high-level interpreter languages, i.e., Python, JavaScript, HTML, and CSS, which is another merit of UNI-EM, i.e., high code maintainability. Although we have provided UNI-EM only on Microsoft Windows 10 (64 bit) UNI-EM can be easily extended to other OSs such as Linux/Mac OS, depending on user request.

There are sophisticated manual segmentation/proofreading tools, such as CATMAID^43^, webKnossos^44^, NeuTu^18^, and Neuroglancer^45^, all of which are target large-scale EM images and require database servers. After conversion and uploading the data to a database server, these tools work efficiently. However, the managements of database servers also require expert knowledge, and upload/download time may be a bottleneck for users that want to run a quick analysis of data of the latest experiment. UNI-EM can handle small to medium-size EM image stacks at a single desktop computer level, which provides an opportunity to test segmentation algorithms before large-scale use involving database servers or cloud systems.

The functions of Dojo are similar to those of VAST lite^24^. VAST is rather a superset of Dojo, because it has more sophisticated paint functions as well as memory management system. VAST can also work for external databases. We have decided not to incorporate it in UNI-EM because VAST is written in C++, and its source code is not open to public. In future, we might integrate VAST by launching a data server on UNI-EM.

In 2D CNN-based segmentation, a 2D CNN produces 2D segmentation images at first, and then these segments are connected across Z slices to obtain 3D objects. UNI-EM only provides 3D labeling and 3D watershed to connect the 2D segments. There are a variety of high-performance Z slice connectors, such as rule-based connectors^15^, multicut algorithms^46^, and the graph-based active learning of agglomeration (GALA)^47^. These connectors should be incorporated into UNI-EM in the future because they may improve not only neuron segmentation, but also the segmentation of other types of objects.

A drawback of DNN-based segmentation is the tedious labor for generating ground truth. Very recently, domain adaptation has been applied to neuron segmentation^48-50^. This is a technique to transfer the ground truth for a type of EM images to that for another type of EM images. Such new technologies may drastically decrease the required amount of manual annotations for new EM datasets. Incorporation of these new technologies into UNI-EM is another important future direction.

## Methods

### RAND score

We utilized foreground-restricted rand scoring (RAND score) as a metric of segmentation performance^8^, which is defined as follows. Suppose *p*_*ij*_ as the joint probability that a target pixel belongs to the object *i* of inferred segmentation and the object *j* of ground truth segmentation (Σ_*ij*_ *p*_*ij*_ = 1). Then, *s*_*i*_=Σ_*j*_ *p*_*ij*_ is the marginal probability for the inferred segmentation, and *t*_*j*_=Σ_*i*_ *p* _*ij*_ is the marginal probability for the ground truth segmentation. Then, the RAND score, 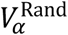, is defined by:

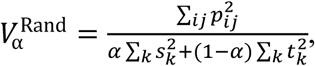

where the Rand F-score *α* is set to 0.5. The split score (α → 0) can be interpreted as precision in the classification of pixel pairs as belonging to the same objects (positive class) or different objects (negative class). The merge score (α → 1) can be interpreted as recall. Generally, 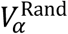 becomes 1 if the segmentation is accurate. This metric was used in ISBI 2012 ^8^.

## Acknowledgements

We would like to thank Ryoji Miyamoto, Noboru Yamaguchi, Hiroko Kita, and Yoshihisa Fujita for their technical assistances. This work was supported partly by the Brain Mapping by Integrated Neurotechnologies for Disease Studies (Brain/MINDS) from the Japanese Agency for Medical Research and Development (AMED), JST CREST (JPMJCR1652), and JSPS KAKENHI (17K00404 to HU and 17H06310 to SI).

## Author contributions

H.U. and S.I. designed the study. H.U wrote all the software programs. T.B. wrote the 2D CNN-based segmentation programs. Y.K. tested the software package and provided an example workflow. H.U., T.B., Y.K., S.O., and S.I. wrote the manuscript.

## References

1 Briggman, K. L. & Bock, D. D. Volume electron microscopy for neuronal circuit reconstruction. Curr Opin Neurobiol 22, 154–161, doi:10.1016/j.conb.2011.10.022 (2012).

2 Helmstaedter, M. Cellular-resolution connectomics: challenges of dense neural circuit reconstruction. Nat Methods 10, 501–507, doi:10.1038/nmeth.2476 (2013).

3 Morgan, J. L. & Lichtman, J. W. Why not connectomics? Nat Methods 10, 494–500, doi:10.1038/nmeth.2480 (2013).

4 Lee, W. C. et al. Anatomy and function of an excitatory network in the visual cortex. Nature 532, 370–374, doi:10.1038/nature17192 (2016).

5 Zheng, Z. et al. A Complete Electron Microscopy Volume of the Brain of Adult Drosophila melanogaster. Cell 174, 730–743 e722, doi:10.1016/j.cell.2018.06.019 (2018).

6 Hildebrand, D. G. C. et al. Whole-brain serial-section electron microscopy in larval zebrafish. Nature 545, 345–349, doi:10.1038/nature22356 (2017).

7 Arganda-Carreras, I., Seung, S., Cardona, A. & Schindelin, J. ISBI Challenge: Segmentation of neuronal structures in EM stacks, http://brainiac2.mit.edu/isbi_challenge/ (2012).

8 Arganda-Carreras, I. et al. Crowdsourcing the creation of image segmentation algorithms for connectomics. Front Neuroanat 9, 142, doi:10.3389/fnana.2015.00142 (2015).

9 Ciresan, D., Giusti, A., Gambardella, L. M. & Jurgen, S. Deep Neural Networks Segment Neuronal Membranes in Electron Microscopy Images. Advances in Neural Information Processing Systems 25, 2843–2851 (2012).

10 Ronneberger, O., Fischer, P. & Brox, T. U-Net: Convolutional Networks for Biomedical Image Segmentation. in Medical Image Computing and Computer-Assisted Intervention—MICCAI 2015 9351, 234–241 (2015).

11 Takemura, S. Y. et al. Synaptic circuits and their variations within different columns in the visual system of Drosophila. Proc Natl Acad Sci U S A 112, 13711–13716, doi:10.1073/pnas.1509820112 (2015).

12 Berning, M., Boergens, K. M. & Helmstaedter, M. SegEM: Efficient Image Analysis for High-Resolution Connectomics. Neuron 87, 1193–1206, doi:10.1016/j.neuron.2015.09.003 (2015).

13 Lee, K., Zung, J., Li, P., Jain, V. & Seung, H. S. Superhuman accuracy on the SNEMI3D connectomics challenge. arXiv preprint, arXiv:1706.00120 (2017).

14 Januszewski, M. et al. High-precision automated reconstruction of neurons with flood-filling networks. Nat Methods 15, 605–610, doi:10.1038/s41592-018-0049-4 (2018).

15 Kaynig, V. et al. Large-scale automatic reconstruction of neuronal processes from electron microscopy images. Med Image Anal 22, 77–88, doi:10.1016/j.media.2015.02.001 (2015).

16 Haehn, D. et al. Scalable Interactive Visualization for Connectomics. Informatics 4, doi:10.3390/informatics4030029 (2017).

17 Bae, J. A. et al. Digital Museum of Retinal Ganglion Cells with Dense Anatomy and Physiology. Cell 173, 1293–1306 e1219, doi:10.1016/j.cell.2018.04.040 (2018).

18 Zhao, T., Olbris, D. J., Yu, Y. & Plaza, S. M. NeuTu: Software for Collaborative, Large-Scale, Segmentation-Based Connectome Reconstruction. Front Neural Circuits 12, 101, doi:10.3389/fncir.2018.00101 (2018).

19 Katz, W. T. & Plaza, S. M. DVID: Distributed Versioned Image-Oriented Dataservice. Frontiers in Neural Circuits 13, doi:10.3389/fncir.2019.00005 (2019).

20 Fiala, J. C. Reconstruct: a free editor for serial section microscopy. J Microsc 218, 52–61, doi:10.1111/j.1365-2818.2005.01466.x (2005).

21 Sommer, C., Straehle, C., Kothe, U. & Hamprecht, F. A. Ilastik: Interactive Learning and Segmentation Toolkit. 2011 8th IEEE International Symposium on Biomedical Imaging: From Nano to Macro, 230–233 (2011).

22 Helmstaedter, M., Briggman, K. L. & Denk, W. High-accuracy neurite reconstruction for highthroughput neuroanatomy. Nat Neurosci 14, 1081–1088, doi:10.1038/nn.2868 (2011).

23 Belevich, I., Joensuu, M., Kumar, D., Vihinen, H. & Jokitalo, E. Microscopy Image Browser: A Platform for Segmentation and Analysis of Multidimensional Datasets. PLoS Biol 14, e1002340, doi:10.1371/journal.pbio.1002340 (2016).

24 Berger, D. R., Seung, H. S. & Lichtman, J. W. VAST (Volume Annotation and Segmentation Tool): Efficient Manual and Semi-Automatic Labeling of Large 3D Image Stacks. Front Neural Circuits 12, 88, doi:10.3389/fncir.2018.00088 (2018).

25 Falk, T. et al. U-Net: deep learning for cell counting, detection, and morphometry. Nat Methods 16, 67–70, doi:10.1038/s41592-018-0261-2 (2019).

26 Abadi, M. et al. in Proceedings of the 12th USENIX conference on Operating Systems Design and Implementation 265–283 (USENIX Association, Savannah, GA, USA, 2016).

27 He, K., Zhang, X., Ren, S. & Sun, J. Identity Mappings in Deep Residual Networks. European Conference on Computer Vision, 630–645 (2016).

28 Srivastava, R. K., Greff, K. & Schmidhuber, J. Highway Networks. arXiv preprint, arXiv:1505.00387 (2015).

29 Huang, G., Liu, Z. & Weinberger, K. Q. Densely Connected Convolutional Networks. arXiv preprint, arXiv:1608.06993 (2016).

30 Haehn, D. et al. Design and Evaluation of Interactive Proofreading Tools for Connectomics. Proceedings IEEE SciVis 20, 2466–2475 (2014).

31 StatCounter Global Stats - Browser, O., Search Engine including Mobile Usage Share. statcounter.com (1999).

32 Arganda-Carreras, I., Seung, H. S., Vishwanathan, A. & Berger, D. R. SNEMI3D: 3D Segmentation of neurites in EM images. (2013).

33 Kasthuri, N. et al. Saturated Reconstruction of a Volume of Neocortex. Cell 162, 648–661, doi:10.1016/j.cell.2015.06.054 (2015).

34 Saxton, W. M. & Hollenbeck, P. J. The axonal transport of mitochondria. J Cell Sci 125, 2095–2104, doi:10.1242/jcs.053850 (2012).

35 Ohno, N. et al. Mitochondrial immobilization mediated by syntaphilin facilitates survival of demyelinated axons. Proc Natl Acad Sci U S A 111, 9953–9958, doi:10.1073/pnas.1401155111 (2014).

36 Frey, T. G. & Mannella, C. A. The internal structure of mitochondria. Trends Biochem Sci 25, 319–324 (2000).

37 Nunnari, J. & Suomalainen, A. Mitochondria: in sickness and in health. Cell 148, 1145–1159, doi:10.1016/j.cell.2012.02.035 (2012).

38 Oztel, I., Yolcu, G., Ersoy, I., White, T. & Bunyak, F. Mitochondria Segmentation in Electron Microscopy Volumes using Deep Convolutional Neural Network. Ieee Int C Bioinform, 1195–1200 (2017).

39 Xiao, C. et al. Automatic Mitochondria Segmentation for EM Data Using a 3D Supervised Convolutional Network. Front Neuroanat 12, 92, doi:10.3389/fnana.2018.00092 (2018).

40 Lee, K., Zlateski, A., Vishwanathan, A. & Seung, H. S. in Proceedings of the 28th International Conference on Neural Information Processing Systems - Volume 2 3573–3581 (MIT Press, Montreal, Canada, 2015).

41 Zeng, T., Wu, B. & Ji, S. DeepEM3D: approaching human-level performance on 3D anisotropic EM image segmentation. Bioinformatics 33, 2555–2562, doi:10.1093/bioinformatics/btx188 (2017).

42 Cardona, A. et al. TrakEM2 software for neural circuit reconstruction. PLoS One 7, e38011, doi:10.1371/journal.pone.0038011 (2012).

43 Saalfeld, S., Cardona, A., Hartenstein, V. & Tomancak, P. CATMAID: collaborative annotation toolkit for massive amounts of image data. Bioinformatics 25, 1984–1986, doi:10.1093/bioinformatics/btp266 (2009).

44 Boergens, K. M. et al. webKnossos: efficient online 3D data annotation for connectomics. Nat Methods 14, 691–694, doi:10.1038/nmeth.4331 (2017).

45 Neuroglancer: WebGL-based viewer for volumetric data, https://github.com/google/neuroglancer (2016).

46 Beier, T. et al. Multicut brings automated neurite segmentation closer to human performance. Nat Methods 14, 101–102, doi:10.1038/nmeth.4151 (2017).

47 Nunez-Iglesias, J., Kennedy, R., Plaza, S. M., Chakraborty, A. & Katz, W. T. Graph-based active learning of agglomeration (GALA): a Python library to segment 2D and 3D neuroimages. Front Neuroinform 8, 34, doi:10.3389/fninf.2014.00034 (2014).

48 Bermudez-Chancon, R., Marquez-Neila, P., Salzmann, M. & Fua, P. A Domain-Adaptive Two-Stream U-Net for Electron Microscopy Image Segmentation. I S Biomed Imaging, 400–404 (2018).

49 Roels, J., Hennies, J., Saeys, Y., Philips, W. & Kreshuk, A. Domain Adaptive Segmentation in Volume Electron Microscopy Imaging. arXiv preprint, arXiv:1810.09734 (2018).

50 Januszewski, M. & Jain, V. Segmentation-Enhanced CycleGAN. bioRxiv preprint, http://dx.doi.org/10.1101/548081 (2019).

